# PKD3 localizes to late endosomes to maintain Rab7-dependent endolysosomal homeostasis

**DOI:** 10.1101/2025.01.10.632486

**Authors:** Elena Gutiérrez-Galindo, Katharina Jursik, Yannick Frey, Florian Meyer, Angelika Hausser

## Abstract

Protein kinase D3 (PKD3) is an important regulator of triple-negative breast cancer (TNBC) progression by promoting invasion, proliferation, and stem cell maintenance. However, the mechanism underlying these cellular functions has remained unclear. Here, we report that endogenous PKD3 localizes to Rab7/Lamp1-positive vesicles in MDA-MB-231 cells cultured on stiff matrices. Notably, upon PKD3 depletion the size of Rab7-positive vesicles is smaller. This correlates with impaired endosomal acidification, which is associated with dysregulated Wnt signaling and a decline in stemness. Our data thus unveil a previously unrecognized role of PKD3 in regulating endolysosomal dynamics that contributes to the maintenance of the cancer stem cell population.

## Introduction

The serine/threonine Protein Kinase D family (PKD, comprising PKD1, PKD2 and PKD3) is involved in several cellular processes including cell migration (Peterburs et al., 2009; Christoforides et al., 2012), proliferation (Karam et al., 2012; Huck et al., 2013; Liu et al., 2020), protein transport (Liljedahl et al., 2001; Yeaman et al., 2004; Bossard et al., 2007; Eisler et al., 2018) secretion (Gali et al., 2024; Eiseler et al., 2016) and EMT (Du et al., 2010), and thus has a key role in tumor progression.

PKD exists as an inhibitory dimer in the cytosol and is activated when bound to endomembranes in a diacylglycerol (DAG)-dependent manner (Reinhardt et al., 2023). Consistent with its localization to endomembranes, numerous studies confirm a function of the PKD family in regulating membrane trafficking (reviewed in Gutiérrez-Galindo et al., 2023), particularly in the secretory pathway, and assign it a major function at the trans-Golgi-network (TGN) (Liljedahl et al., 2001; Hausser et al., 2005; Eisler et al., 2018). However, while it has been speculated that PKD2 and PKD3 isoforms can form homo- and heterodimers to control secretion at the level of the Golgi (Bossard et al., 2007), little is known about the spatio-temporal regulation and localization of the different isoforms and how this affects PKD-dependent functions at the endogenous level.

In triple-negative breast cancer (TNBC), PKD2 and PKD3 are expressed with PKD3 being the predominant isoform (Borges et al., 2015; Huck et al., 2013). Recently, an important role for PKD2 in regulating the pro-invasive secretome of TNBC cells has been described (Gali et al., 2024), while PKD3 was shown to be essential to maintain the stem cell population in TNBC cells in both *in vitro* and *in vivo* (Lieb et al., 2020). However, the mechanism by which PKD3 controls this process remained unknown.

Here, we provide evidence that endogenous PKD3 localizes to membranes of the endolysosomal compartment to control Rab7 localization and, as a consequence, retromer complex distribution and endolysosomal acidification. Additionally, we found that this role of PKD3 in preserving endolysosomal homeostasis and function supports stemness in TNBC via modulating the Wnt pathway.

## Results and discussion

### Active PKD3 localizes to membranes of the endolysosomal compartment

To study the molecular role of PKD3 in TNBC cells, we generated MDA-MB-231 cells overexpressing PKD3-GFP in a doxycycline (Dox)-inducible manner. Unforeseen, active PKD3, visualized by staining of the phosphorylated activation loop (S731/735, pPKD), was not detected at a perinuclear compartment but rather localized to vesicles (Figure 1A). However, since this dominant vesicular localization of active PKD3 was only observed in a few cells, we moved to a more physiological 3D-on top culture system that resembles the stiffening of the tumor occurring during BC progression (Levental et al., 2009; Wei et al., 2015; Fattet et al., 2020). When cells were cultured on 5 kPa polyacrylamide (PAA) hydrogels, significantly more cells contained active, vesicle-localized PKD3 in comparison to a standard 2D culture, and the average number of vesicles with active PKD3 per cell was significantly higher (Figure 1B-C). Consequently, all further experiments with MDA-MB-231 cells were performed using the 3D-on top culture system, which appears to favor elevated vesicular PKD3 activity.

**Figure 1.**
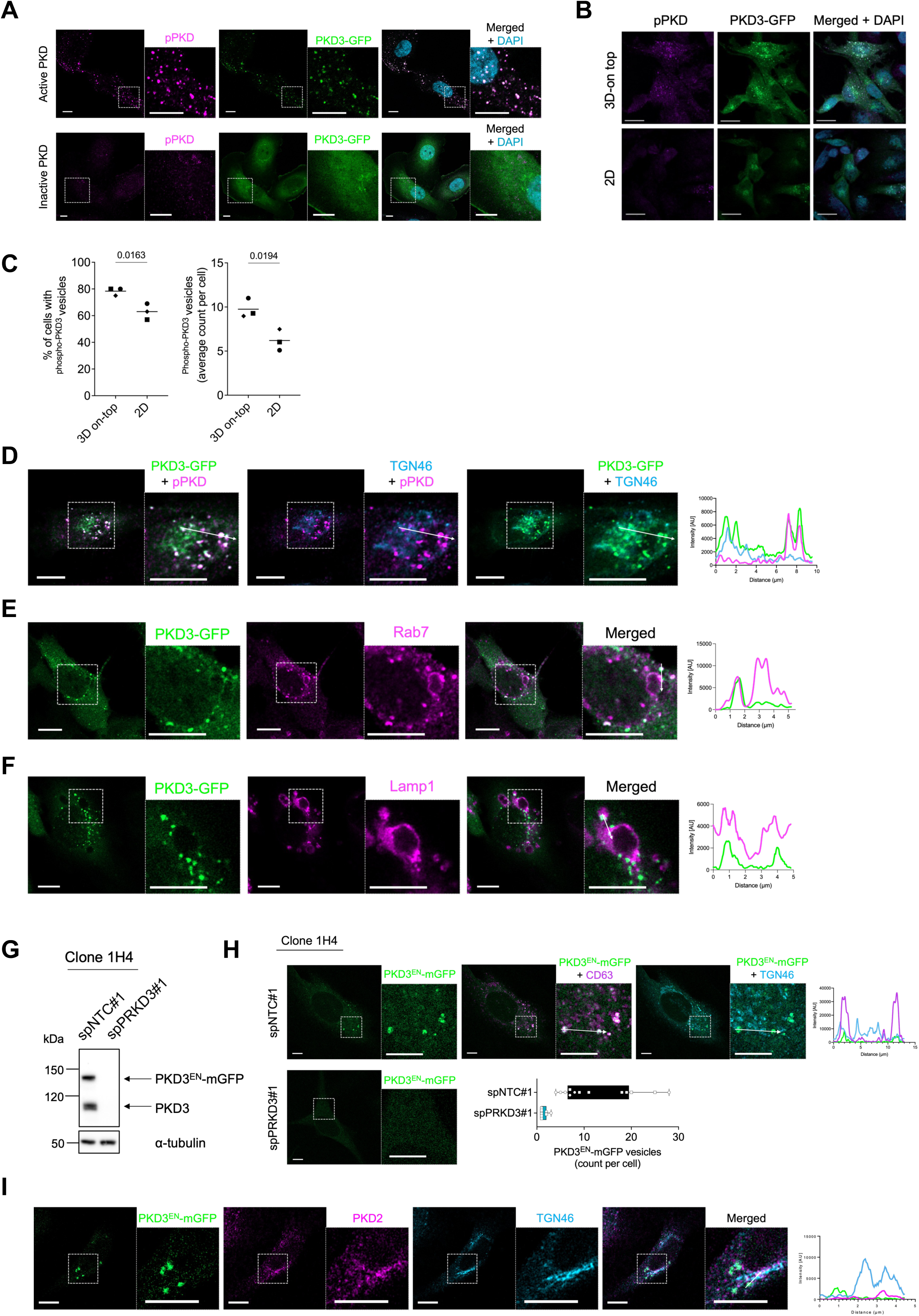
Active PKD3 localizes to membranes of the endolysosomal compartment. **A)** Maximum intensity projections (MIP) of MDA-MB-231_PKD3-GFP cells seeded on glass coverslips, fixed and stained for pPKD (magenta) and DAPI (blue); scale bar: 10 μm. **B)** MIPs of MDA-MB-231_PKD3-GFP cells seeded on glass coverslips (2D) or 5 kPa PAA hydrogels (3D-on top), fixed and stained for pPKD (magenta) and DAPI (blue); scale bar: 20 μm. **C)** Quantification of B); n= 3, N > 35 cells per condition and experiment; statistical comparison by unpaired t-test. **D-F)** MDA-MB-231_PKD3-GFP cells were seeded on 5 kPa PAA hydrogels, fixed and stained for pPKD (magenta) and TGN46 (blue) in D), Rab7 (magenta) in E) and Lamp1 (magenta) in F); a middle Z-section is shown; scale bar: 10 µm; the histogram represents the intensity profile in the area marked with a white arrow in the merged image. **G)** Whole cell lysates of HeLa_ PKD3^EN^-mGFP (Clone 1H4) transiently transfected with spNTC#1 or spPRKD3#1 were subjected to Western blot analysis and probed for PKD3; α-tubulin was used as a loading control. **H)** HeLa_ PKD3^EN^-mGFP cells (Clone 1H4) transiently transfected with spNTC#1 or spPRKD3#1 were seeded on glass coverslips and fixed; control samples were stained for CD63 (magenta) and TGN46 (blue); scale bar: 10 µm; the histogram represents the intensity profile in the area marked with a white arrow; the graph represents the number of PKD3^EN^-mGFP positive vesicles per cell from two independent experiments. **I)** HeLa_ PKD3^EN^-mGFP cells (Clone 1H4) were seeded on glass coverslips, fixed and stained for PKD2 (magenta) and TGN46 (blue); a middle Z-section is shown; scale bar: 10 µm; the histogram represents the intensity profile in the area marked with a white arrow in the merged image.

PKD is active when bound to endomembranes (Hausser et al., 2002) and, in epithelial cells, the ectopically expressed protein localizes to the TGN to control the fission of vesicles containing cargo for the plasma membrane (Liljedahl et al., 2001; Hausser et al., 2005; Eisler et al., 2018). Breast cancer cell lines often harbor a fragmented Golgi apparatus (Sewell et al., 2006). Hence, to investigate whether the vesicles on which PKD3-GFP was detected might originate from the TGN, we stained PKD3-GFP expressing MDA-MB-231 cells with an antibody specific for TGN46. To our surprise, PKD3-GFP was frequently localized near perinuclear, TGN46 positive membranes, but did not co-localize with them. Furthermore, the vesicles positive for active PKD3-GFP (pPKD), also did not colocalize with TGN46 (Figure 1D). We therefore tested different markers for endosomal membranes and found that PKD3-GFP vesicles co-localized with Rab7 (Figure 1E), Lamp1 (Figure 1F), and CD63 (Figure S1A), all well-established components of late endosomal and lysosomal membranes (Van Der Beek et al., 2021). In line with this, active PKD3-GFP was often found to decorate the membrane of acidic vesicles labelled with LysoTracker (Figure S1B).

To rule out that this localization is a result of the ectopic overexpression of PKD3-GFP, we used CRISPR/Cas12a assisted PCR tagging (Fueller et al., 2020) to fuse endogenous PKD3 (PKD3^EN^) with mGFP. Since our tagging approach was not successful in MDA-MB-231 cells, we employed the well-established HeLa cells that express PKD2 and PKD3, but not PKD1 (Yeaman et al., 2004; Döppler et al., 2014), as a model. In brief, after transfection and antibiotic selection to generate two independent HeLa_PKD3^EN^-mGFP pools, PKD3^EN^-mGFP expression was controlled by Western blot (Figure S1C), and a subsequent limited dilution of pool 2 was performed to select positive single clones. Out of 32 clones, 7 were positive for PKD3^EN^-mGFP expression and clone 1H4 and 2G2 were chosen for further experiments (Figure S1D). Of note, all clones only showed a partial tagging efficiency since expression of untagged PKD3 was still detectable. Sequencing of the genomic DNA of both clones showed the correct in frame insertion of mGFP at the C-terminus of PKD3 (Figure S1E). In addition, transient transfection with a PKD3 specific siRNA (#spPRKD3#1) proved specificity as PKD3^EN^-mGFP levels decreased together with untagged PKD3 for Pool 2 (Figure S1C) and both clones, 1H4 (Figure 1G) and 2G2 (Figure S1F). Confocal microscopy studies of the two clones revealed a vesicular localization of PKD3^EN^-mGFP that was absent in the knockdown condition (Figure 1H, S1G). More precisely, these PKD3^EN^-mGFP vesicles co-localized with CD63, and did not show any co-localization with the perinuclear TGN (detected with TGN46), for both 1H4 (Figure 1H) and 2G2 clones (Figure S1G). As previously described for MDA-MB-231_PKD3-GFP cells, these PKD3^EN^-mGFP positive, late endosomal membranes were also positive for Rab7 (Figure S1H) confirming our findings.

The formation of homo-and heterodimers for all PKD isoforms has been reported in different studies using recombinant proteins (Bossard et al., 2007; Aicart-Ramos et al., 2016; Elsner et al., 2019), but whether these are formed at the endogenous level still remains unclear. Since HeLa cells express PKD2 but not PKD1, we stained endogenous PKD2 in HeLa_PKD3^EN^-mGFP cells using a specific antibody. We found a co-localization of PKD2 with TGN46, but no evident overlap with PKD3^EN^-mGFP (Figure 1I). This suggests that the majority of endogenous PKD2 and PKD3 exist as separate entities rather than forming heterodimers under basal conditions.

Taken together, our observations indicate a previously unrecognized localization of endogenous PKD3 to membranes of the endolysosomal compartment.

### Loss of PKD3 leads to a redistribution of Rab7

To further explore the role of PKD3 at endolysosomes, we investigated possible binding partners of the kinase at endolysosomes. Two independent publications detected by proximity label-mass spectrometry an interaction of PKD3 with Rab7A (hereafter Rab7) (Buljan et al., 2020; Yan et al., 2022), which is well known to be the key regulator of early-to-late endosome maturation and motility, lysosomal biogenesis and cargo transport (Guerra and Bucci, 2016) and which we found decorating PKD3-GFP positive vesicles (Figure 1E, S1H).

To investigate whether PKD3 can be functionally linked to Rab7, MDA-MB-231 cells transiently transfected with a control siRNA (spNTC#1), or a PKD3-specific siRNA (spPRKD3#1) were stained for Rab7, along with the lysosomal proteins Lamp1 or CD63. In control cells, Rab7 was enriched at vesicular membranes partly overlapping with Lamp1/CD63 (Figure 2A). Depletion of PKD3 reduced Rab7 localization to Lamp1-/CD63-positive membranes (Figure 2B) and Rab7 appeared to be more distributed throughout the cell (Figure 2A). Quantification of the Rab7 signal showed a decrease of the mean fluorescence intensity (MFI) in PKD3 depleted cells compared to control cells (Figure 2C). Additionally, the Rab7-positive vesicles were analysed to compare size and count, demonstrating a shift from larger to smaller vesicles upon PKD3 depletion (Figure 2D), while the area of Lamp1-/CD63-positive vesicles remained unchanged (Figure 2E). Western blot analysis confirmed equal expression of Rab7 in all conditions analysed, supporting that the differences in fluorescence intensity are due to the redistribution of Rab7 rather than loss of the protein (Figure 2F). Notably, this redistribution was not accompanied by a detectable change in Rab7 activity, as GST-RILP pulldown assays showed no differences between control and PKD3-depleted cells (Figure 2G), ruling out a cytosolic localization of Rab7. While Rab7 is predominantly localized to late endosomes and lysosomes, our findings align with previous reports indicating that Rab7 can also associate with small vesicles, the nature of which remains to be determined (Rojas et al., 2008; Van Der Beek et al., 2021).

**Figure 2.**
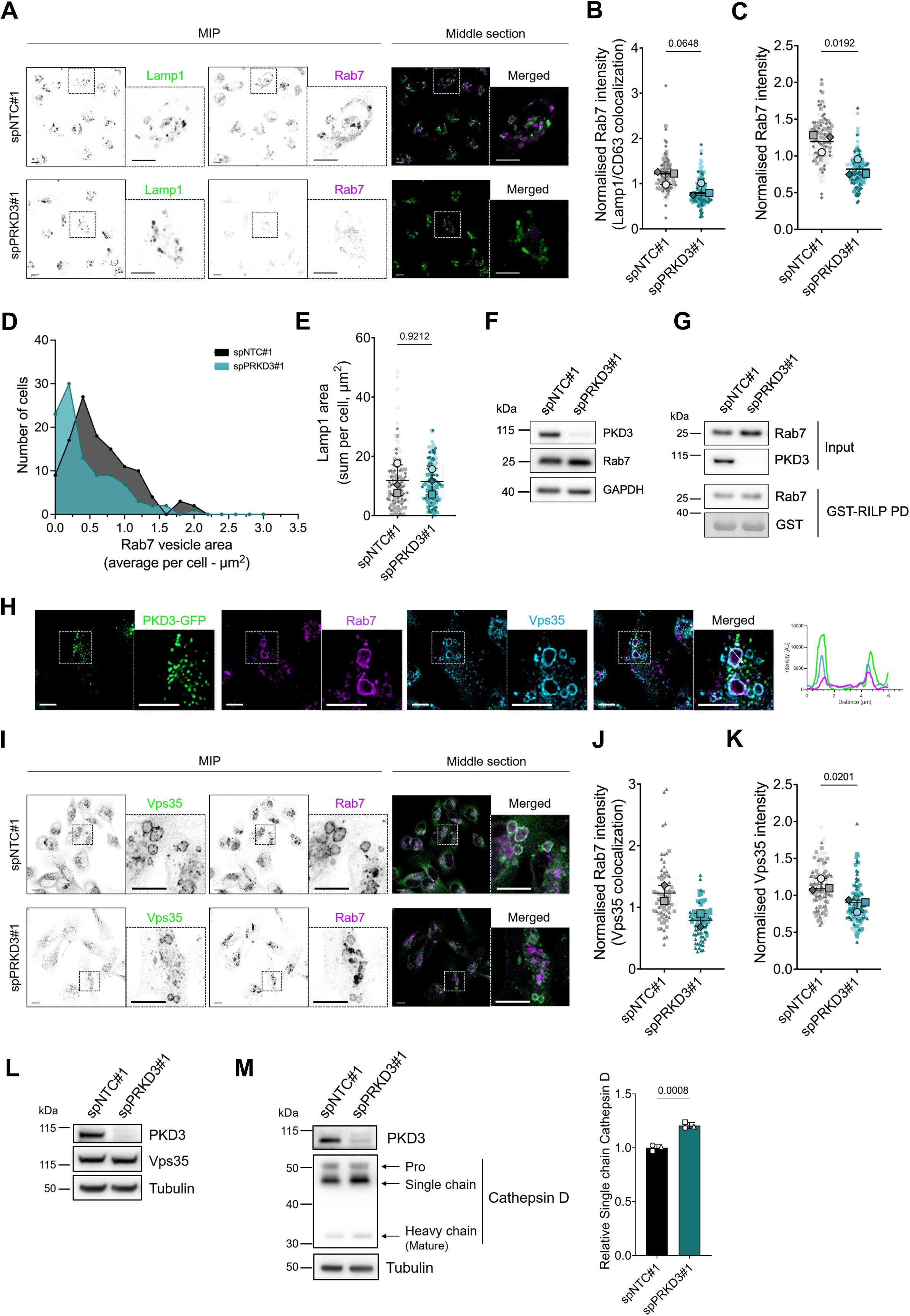
Loss of PKD3 leads to a redistribution of Rab7. **A)** MDA-MB-231 cells transiently transfected with spNTC#1 or spPRKD3#1 were seeded on 5 kPa PAA hydrogels, fixed and stained as indicated: Lamp1 (green) and Rab7 (magenta); black and white images show maximum intensity projections (MIP), and colored images show a middle Z-section; scale bar: 10 µm. **B-E)** Quantification of A); n=3; N>35 cells per condition and experiment. **B)** Colocalization of Rab7 with Lamp1 or CD63, analised as the MFI of Rab7 on Lamp1- or CD63-positive vesicles; statistical comparison by unpaired t-test. **C)** Rab7 mean fluorescence intensity (MFI) of MIPs, normalised to the average MFI for each independent experiment; statistical comparison by unpaired t-test. **D)** Frequency of distribution of Rab7-positive vesicles from three independent experiments, distributed by area. **E)** Area of Lamp1- or CD63-positive vesicles (sum per cell); statistical comparison by unpaired t-test. **F)** Whole cell lysates of MDA-MB-231 cells transiently transfected with spNTC#1 or spPRKD3#1 were subjected to Western blot analysis and probed for PKD3 and Rab7; GAPDH was used as a loading control. The blot is representative of three independent experiments. **G)** Whole cell lysates of MDA-MB-231 cells transiently transfected with spNTC#1 or spPRKD3#1 were subjected to pull-down assays using GST-RILP and analysed by Western blot using a Rab7 antibody. The blot is representative of three independent experiments. **H)** MDA-MB-231_PKD3-GFP cells were seeded on 5 kPa PAA hydrogels for, fixed and stained for Rab7 (magenta) and Vps35 (blue); a middle Z-section is shown; scale bar: 10 µm; the histogram represents the intensity profile in the area marked with a white arrow in the merged image. **I)** MDA-MB-231 cells transiently transfected with spNTC#1 or spPRKD3#1 were seeded on 5 kPa PAA hydrogels, fixed and stained as indicated: Vps35 (green) and Rab7 (magenta); black and white images show MIPs, and colored images show a middle Z-section; scale bar: 10 µm. **J-K)** Quantification of I). **J)** Colocalization of Rab7 and Vps35, analised as the MFI of Rab7 on Vps35-positive vesicles; n=2, N>30 cell per condition and experiment. **K)** Vps35 MFI of MIPs, normalised to the average MFI for each independent experiment; statistical comparison by unpaired t-test; n=3, N>30 cell per condition and experiment**. L-M)** Whole cell lysates of MDA-MB-231 cells transiently transfected with spNTC#1 or spPRKD3#1 were subjected to Western blot analysis and probed for: L) PKD3 and Vps35; M) PKD3 and CathepsinD; α-tubulin was used as a loading control. The blots are representative of three independent experiments. The graph shows the relative levels of single chain CathepsinD; statistical comparison by t-test; n=3.

Previous studies have demonstrated that Rab7 localization to late endosomal membranes is associated with the recruitment of the retromer complex (Rojas et al., 2008; Seaman et al., 2009). This multiprotein complex regulates the sorting of integral membrane proteins through the endolysosomal network and plays a critical role in Rab7 nucleotide cycling (Seaman, 2012; Burd and Cullen, 2014; Jimenez-Orgaz et al., 2018; Kvainickas et al., 2019). Remarkably, we observed that PKD3-GFP decorated the membrane of enlarged Rab7-positive vesicles, where the retromer subunit Vps35 was also found (Figure 2H). Furthermore, since PKD3 expression modulated the enrichment of Rab7 at endosomal membranes and Rab7 is closely connected with the retromer complex, we hypothesized that the recruitment of the retromer complex to these membranes could follow a similar pattern. Indeed, co-staining of Rab7 along with the retromer subunit Vps35 in MDA-MB-231 cells confirmed a reduced co-localization of both proteins when PKD3 was depleted (Figure 2I-J). Quantification of the Vps35 signal also showed an overall reduced recruitment of retromer to endomembranes (Figure 2K), while Vps35 protein levels remained unchanged (Figure 2L). These findings suggest that the retromer function could be potentially modulated by endolysosomal PKD3. A prominent cargo of the retromer complex are the lysosomal cathepsins (Rojas et al., 2008; Cui et al., 2019), whose proper trafficking and maturation along the endolysosomal pathway is Rab7 dependent (Rojas et al., 2008). We therefore analysed the maturation of cathepsin D (CatD) by Western blot analysis. In control cells, three protein bands were detected, presenting pro-CatD (52 kDa), an active intermediate single chain CatD (48 kDa) and the mature heavy chain (34 kDa) (Mijanovic et al., 2021). Notably, in PKD3 depleted cells, the 48 kDa form, presenting endosome localized single-chain CatD, was significantly enriched compared to control cells (Figure 2M) suggesting deregulated trafficking of CatD at the endolysosomal compartment. Our result is in line with previous findings (Huck et al., 2013) and proposes a role for PKD3 in coordinating endolysosomal trafficking and consequently lysosomal function via Rab7.

### PKD3 supports endolysosomal acidification to regulate the Wnt pathway

In TNBC cells, PKD3 has been functionally linked to the maintenance of the stem cell population (Lieb et al., 2020). Likewise, the Wnt/β-catenin signaling pathway, which is one of the major contributors in maintaining stemness (Zhang et al., 2023) is elevated in breast cancer stem cells (Jang et al., 2015). Lysosomal function is closely linked to stem cell maintenance in breast cancer (Choi et al., 2014; Turcu et al., 2020; Nowosad and Besson, 2022) while Wnt signaling has been linked to an increase in the acidity of the endolysosomal compartment and lysosomal activity (Azoulay-Alfaguter et al., 2014; Albrecht et al., 2020; Avrahami et al., 2020). These connections suggest that our findings on PKD3 localization and its role in endolysosomal function may critically influence Wnt signaling and, consequently, cancer stemness.

To address this issue, we first performed a sphere formation assay (SFA) and confirmed that under our 3D-on-top culture conditions, the sphere formation efficiency (SFE) decreased with PKD3 depletion (Figure S2A), confirming previous results (Lieb et al., 2020). Next, we investigated a potential role of PKD3 in the Wnt pathway and employed the GSK3β inhibitor CHIR99021 to induce canonical Wnt signaling in MDA-MB-231 cells (Albrecht et al., 2020). As a readout for Wnt activity, we detected non-phosphorylated β-catenin, one of the main effector proteins in the canonical Wnt signaling (reviewed in (Verheyen and Gottardi, 2010). Indeed, in control cells, CHIR99021 induced a potent activation of β-catenin visible by the enrichment of the non-phosphorylated protein. Upon PKD3 depletion, either by transient transfection of a PKD3-specific siRNA (Figure 3A) or by a stable knockdown (shPKD3#2, Figure 3B), the amount of active β-catenin was significantly reduced compared to control cells. Likewise, when PKD activity was inhibited using the PKD pan-inhibitor CRT0066101 prior treatment with CHIR99021, a decrease in non-phosphorylated β-catenin was observed (Figure 3C). To further corroborate these data, we conducted qPCR analysis of *Axin2*, widely considered a marker for the Wnt signaling pathway in TNBC (MacDonald et al., 2009; Maubant et al., 2015). Depletion of PKD3 either by transient transfection of PKD3-specific siRNAs or by stably expressed shPRKD3#2, reduced the mRNA levels of *Axin2* in comparison to the control conditions (Figure 3D-E). Additionally, both MDA-MB-231 and BT549, another TNBC cell line, showed lower expression of *Axin2* when PKD activity was inhibited by CRT0066101 treatment (Figure S2B). Taken together, these results demonstrate an involvement of PKD3 in the Wnt/β-catenin signaling pathway in TNBC cells.

**Figure 3.**
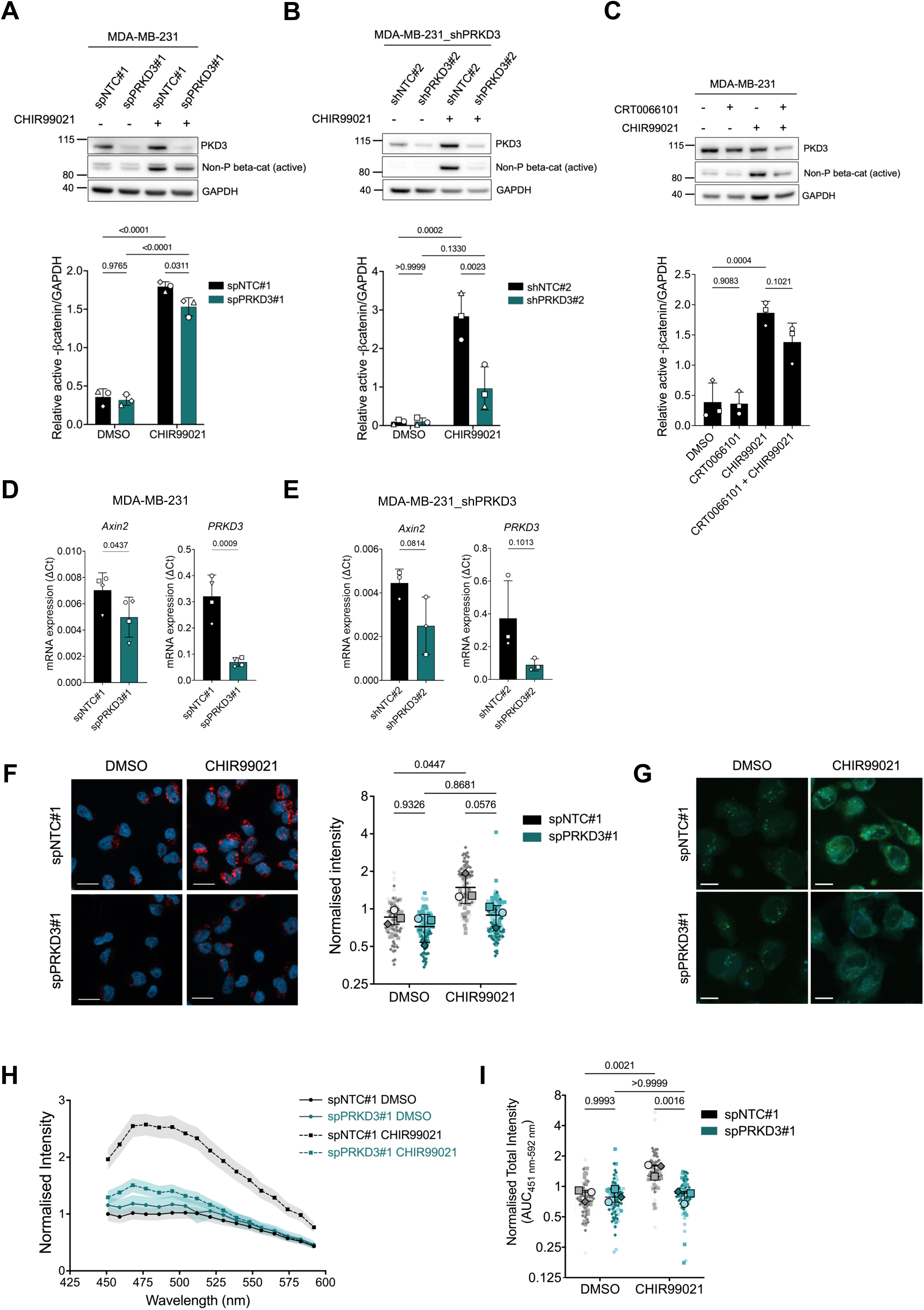
PKD3 supports lysosomal acidification to regulate the Wnt pathway. **A-C)** Western blot analysis of cells seeded on 5 kPa PAA hydrogels, treated as indicated: MDA-MB-231 transiently transfected with spNTC#1 or spPRKD3# (A) or stably expressing shNTC#2 or shPRKD3#2 (B) were treated with DMSO or CHIR99021 (8 nM) for 2 h; C) MDA-MB-231 cells were treated with DMSO or CRT0066101 (2.5 µM) for 1 h prior treatment with DMSO or CHIR99021 (8 nM) for 2 h; whole cell lysates were subjected to Western blot analysis and probed for PKD3 and non-phosphorylated β-catenin; GAPDH was used as a loading control; the upper panels show a representative experiment and the graphs below show the relative levels of non-phosphorylated β-catenin in three independent experiments, normalized to GAPDH levels; statistical comparison by two-way ANOVA with Šídák’s multiple comparisons test (A-B) or ordinary one-way ANOVA with Holm-Šídák’s multiple comparisons test (C). **D-E)** qPCR analysis of *Axin1* and *PRKD3* in cells seeded on 5 kPa PAA hydrogels; D) MDA-MB-231 cells were transiently transfected with spNTC#1 or spPRKD3#1; n=4; E) MDA-MB-231 cells stably expressed shNTC#2 or shPRKD3#2; n=3); statistical comparison by t-test. **F-I)** MDA-MB-231 cells transiently transfected with spNTC#1 or spPRKD3#1 were seeded on 5 kPa PAA hydrogels, treated with DMSO or CHIR99021 (8 nM) for 2 h, incubated with LysoTracker (F) or LysoSensor (G) and fixed; F) Lysotracker; scale bar: 20 µm; data in graph represents the MFI, normalized to the average MFI for each independent experiment; statistical comparison by two-way ANOVA with Šídák’s multiple comparisons test; n= 3, N > 20 cells per condition and experiment. H) LysoSensor intensity measured in the 450-600 nm range per vesicle; scale bar: 10 µm; data in graph represent the average acidic vesicle intensity per cell, normalized to the average intensity for each independent experiment; the graph shows a representative experiment, and values are expressed in mean ± 95% CI. I) LysoSensor intensity: Area under the curve (AUC) of three independent experiments; N > 5-30 cells per condition and experiment. Statistical comparison by two-way ANOVA with Šídák’s multiple comparisons test.

Activation of canonical Wnt signaling via inhibition of GSK3β has been previously reported to regulate endosomal luminal pH (Azoulay-Alfaguter et al., 2014; Albrecht et al., 2020; Avrahami et al., 2020) and to correlate with Rab7 localization to lysosomes (Avrahami et al., 2020). In fact, a fast drop in the intraluminal pH has been detected during Rab conversion (Podinovskaia et al., 2021). For this reason, we next investigated whether PKD3 activity is involved in regulating endosomal acidification. To this end, control and PKD3-depleted MDA-MB-231 (Figure 3F) and BT549 (Figure S2C) cells were left untreated or treated with CHIR99021 as previously described and incubated with LysoTracker Red. Confocal imaging studies showed a strong and significant increase in LysoTracker dye intensity in control cells treated with the GSK3β inhibitor confirming previous findings (Avrahami et al., 2020). Notably, in PKD3-depleted cells, this CHIR99021-induced increase was not visible (Figure 3F, S2C) showing that PKD3 is required for acidification of lysosomes upon induction of Wnt signaling. We did not detect a significant difference under basal conditions, but a trend toward lower LysoTracker dye intensity for PKD3 depletion was observed in both cell lines.

Additionally, we analysed the impact of PKD3 activity on acidic vesicles using LysoSensor Yellow/Blue DND-160, which exhibits dual spectral peaks that are pH-dependent. Acidification of the lysosomes detected with the probe was quantified by measuring the intensity of the signal at 451-592 nm, corresponding to the emission-spectra in more acidic organelles. As expected, CHIR99021 treatment resulted in a strong increase in fluorescence intensity in control cells, indicating a drop in the luminal pH. Importantly, PKD3 depleted cells failed to acidify the lysosomes in response to the treatment, corroborating our LysoTracker experiments (Figure 3G-I). Our data are also in line with recent data showing that the PKD activator and tumor promoter phorbolester increased Wnt-driven colorectal cancer progression, suggesting a requirement of lysosomal acidification for Wnt signaling (Tejeda-Munoz et al., 2023). Since we were unable to excite the probe in the UV range to detect blue fluorescence, a ratiometric measurement to detect organelles with neutral intraluminal pH was not possible. This could explain why we did not observe significant differences in luminal pH in PKD3-depleted cells under basal conditions albeit a trend in the LysoTracker experiments was visible. Nevertheless, with these experiments, we provide evidence that PKD3 is necessary to maintain acidification of endolysosomes in TNBC cells.

### Concluding remarks

Our study provides insights into a previously unrecognized localization of endogenous PKD3 to endolysosomes and assigns the kinase a function in the localization of Rab7 to these membranes. Loss of PKD3 leads to a redistribution of Rab7, which affects the localization of the retromer complex and proper lysosomal function.

In our studies, using 3D-on top culture seemed to be key to identify active PKD3 at endolysosomal vesicles, while its basal activity using standard 2D culture was rather low. This could explain why other studies showed only a few cells where PKD3 was identified in punctae, and these were therefore not fully characterized (Lu et al., 2007; Huck et al., 2013). We further demonstrate that endogenous PKD2 localizes to the TGN and does not overlap with endogenous tagged PKD3^EN^-mGFP, supporting an isoform-specific function for PKD3 homodimers at endolysosomes. Notably, PKD2 and PKD3 differ in structure, with PKD3 lacking the postsynaptic density protein-95/disclarge tumor suppressor protein/zonula occludens-1 (PDZ) motif (Sánchez-Ruiloba et al., 2006) and both kinases possessing distinct intrinsically disordered regions (Gutiérrez-Galindo et al., 2023b). These differences could control specific protein-protein interactions, which in turn, together with DAG, determine the localization of the kinases. Our findings are further in line with a recent study demonstrating a predominant role for PKD2 in regulating the PKD-dependent pro-invasive secretome in metastatic TNBC cells, with only a minor contribution of PKD3 (Gali et al., 2024).

Furthermore, the here reported defect in acidification upon PKD3 depletion affects the Wnt signaling pathway. Remarkably, *PRKD3* was among the genes up-regulated in Wnt3a-stimulated TNBC cells (Maubant et al., 2015), further substantiating a role for the kinase in Wnt signaling. Likewise, by combining proteomics and transcriptomics, Muñoz et al. defined the stem cell signature in the intestine, where the Wnt pathway is one of the major forces controlling homeostasis (Krausova and Korinek, 2014), and which included PKD3 among other kinases (Muñoz et al., 2012). These studies support our hypothesis for a critical role of PKD3 in the Wnt signaling pathway through regulating Rab7 at late endosomes and thus luminal acidification. In line with the PKD3-dependent localization of Vps35, the retromer complex has been also related to Wnt signaling, via recycling Wntless from endosomes to the TGN to sustain Wnt secretion (Belenkaya et al., 2008).

In MDA-MB-231 cells, Rab7 localization to enlarged endosomes was PKD3 dependent. Rab7 controls endolysosomal function via distinct mechanisms and binding different effectors (Guerra and Bucci, 2016; Mulligan and Winckler, 2023). For example, binding to the retromer subunit Vps35 regulates cargo retrieval for hydrolase transport (Rojas et al., 2008; Seaman et al., 2009; Cui et al., 2019), while acidification is dependent on the interaction with RILP and the assembly of the V-ATPase (Mulligan et al., 2024). Our data showing that PKD3 depleted cells display a decrease in Rab7 co-localization with Vps35 and endolysosomal acidification could explain the observed defect in CatD maturation. Indeed, a low pH is necessary for the dissociation of mannose-6-phosphate-receptor (M6PR), as well as for the cleavage and activation of hydrolases (Hu et al., 2015). In any case, the presence of active Rab7 on endolysosomal membranes is indispensable for their proper function (Guerra and Bucci, 2016) and our data place PKD3 upstream of this process.

Within the scope of these studies, a specific PKD3 substrate at endolysosomal membranes has not been identified. Rab7 activity is regulated by the GEF Mon1-Ccz1 (Nordmann et al., 2010) and different GAP proteins with a conserved Tre2/Bub2/Cdc16 (TBC) domain (Stroupe, 2018). Of these, TBC1D5 is of particular interest since it binds with high affinity to the retromer complex at maturing and late endosomes (Seaman et al., 2009, 2018; Jia et al., 2016; Kvainickas et al., 2019). Indeed, recent studies suggest a role for retromer-associated TBC1D5 in regulating the distribution of active Rab7 pools to maintain endolysosomal function (Jimenez-Orgaz et al., 2018; Kvainickas et al., 2019), a role that we show here for PKD3. Our work thus encourages future studies to identify potential PKD3 substrates at endolysosomal membranes, which promote the oncogenic role of lysosomes in TNBC.

## Materials and methods

### Cell lines and reagents

The TNBC cell lines MDA-MB-231, BT549, MDA-MB-231-based knockdown cells (in text: shNTC#2 and shPRKD3#2; (Borges et al., 2015)), MDA-MB-231_PKD3-GFP and HeLa_PKD3^EN^-mGFP (both generated in this study) were cultured in high-glucose Dulbecco’s modified Eagle’s medium (DMEM, Gibco). HeLa cells were cultured in RPMI-1640 (Gibco). All cell lines were supplemented with 10% Fetal Bovine Serum (FBS, Sigma #F7524), cultured at 37°C in a humidified chamber with 5% CO_2_, and maintained until 80-90% confluence prior passaging, for no longer than 2 months. MDA-MB-231_PKD3-GFP cells were maintained with 500 µg/ml G418 sulfate (Geneticin, Calbiochem #345810) and HeLa_PKD3^EN^-mGFP with 2 µg/ml Puromycin (Thermo Fisher Scientific).

MDA-MB-231_PKD3-GFP cells were generated by lentiviral transduction. The PKD3-GFP encoding module was subcloned from pEGFP-N1-PKD3 (Huck et al., 2012) into the vector pCW57-MCS1-P2A-MCS2 (Addgene #89180) by AgeI restriction using the NEBuilder HiFi DNA Assembly Cloning Kit (New England Biolabs #E55220). Lentiviral particles were generated by cotransfecting LentiX HEK293T cells (provided by Dr. Philipp Rathert, University of Stuttgart, and maintained in DMEM with 10% FBS) with virus packaging vectors psPAX2 (Addgene #12260), pCMV-V-SVG (Addgene #8454) and pCW57-MCS1-P2A-MCS2-PKD3-GFP. Transfection was performed using a 1:3 (w/w) mixture of DNA to polyethylenimine (Sigma Aldrich). The resulting virus soup was collected and filtered 24 h after transfection and used for transduction of MDA-MB-231 cells in combination with 8 µg/ml polybrene (Millipore). The transfected cells were selected with 500 µg/ml G418 sulfate. The successful expression of PKD3-GFP was assessed by immunofluorescence and Western blot. During the experiments, PKD3-GFP expression was induced with 1 µg/ml doxycycline for the last 24 h of the experiment.

For reverse transient transfection, 2 x 10^5^ cells were seeded into 6-well plates and transfected with 15 pmol of a mix of four individual siRNAs (Non-Targeting Control or Human *PRKD3*; in text, spNTC#1 or spPRKD3#1; ON-TARGETplus SMARTpool, Horizon Discovery), using Lipofectamine RNAiMAX Transfection Reagent (Invitrogen #13778150) for TNBC cell lines and TransIT-HeLaMONSTER Transfection Kit (Mirus Bio #MIR2904) for HeLa and HeLa_PKD3^EN^-mGFP cells.

### CRISPR/Cas12a assisted PCR tagging

Endogenous PKD3 was tagged at the C-terminus using a PCR-based CRISPR-Cas12a approach (Fueller et al., 2020). A modified plasmid pMaCTag-P05 (Addgene #120016), harboring the monomerizing A206K mutation in EGFP, was used as PCR template with the oligos M1_PKD3: 5′-cgc tgg gaa ata cat gca tac aca cat aac ctt gta tac cca aag cac ttc att atg gct cct aat cca gat gat atg gaa gaa gat cct tca ggt gga gga ggt agt g-3′ and M2_PKD3: 5′-agc aaa ata tca gtc cat aaa atg aaa tcc ttc ctt att tag gtt agc tca gtg aaa aaa aag tga tta agg atc ttc ttc atc tac aag agt aga aat tag cta gct gca tcg gta cc-3′. Cells were transfected with purified PCR product and enAsCas12a helper plasmid (a gift from Keith Joung and Benjamin Kleinstiver; Addgene #107941; (Kleinstiver et al., 2019)) using Lipofectamine 2000 (Thermo Fisher #11668019). Three days after transfection puromycin was added at a concentration of 2 µg/ml.

### Genomic DNA isolation and PCR

Genomic DNA was isolated from HeLa_PKD3^EN^-mGFP cells using the PureLink Genomic DNA Kit (Thermo Fisher #K182001) according to the manufacturer’s instruction. The C-terminal region of the *PRKD3* gene was amplified from genomic DNA by PCR using the primers GRCh38.p1471479 for (5′-cct agg act atc aga cttg gct-3′) and GFP SalI rev (5′-aaa gtc gac gcc ttg tac agc tcg tcc atg cc-3′). After purification, the PCR product was sequenced using the primer GRCh38.p1471479 for.

### Preparation of polyacrylamide (PAA) hydrogels

Polyacrylamide (PAA) hydrogels were prepared based on the protocol previously described (Tse and Engler, 2010). Briefly, 18 or 24 mm square No. 1.5, or 42 mm circle coverslips were incubated with 0.1 M NaOH for 5 minutes, dried, incubated with 3-Aminopropyltriethoxysilane (APTES, Sigma #440140) for 3 minutes, washed with ddH_2_O for 30 minutes, dried, incubated with 0.5% glutaraldehyde (Sigma #G7651) in PBS for 30 minutes and dried. A mix of acrylamide and bis-acrylamide (BioRad #1610140, #1610142) was prepared according to desired stiffness (5 kPa) based on the publication above and polymerized with ammonium persulfate (APS) and Tetramethylethylenediamine (TEMED, Roth #2367). The mixture was polymerized between the amino-silanated coverslip and a Dichlorodimethylsilane (DCDMS, PlusOne GE Healthcare #17-1332-01)-coated coverslip for 15 minutes. After removing the top coverslip, the hydrogels were rinsed twice with PBS and stored at 4°C. Prior use, hydrogels were functionalized with 0.2 mg/ml sulfo-SANPAH (Thermo Scientific, #22589) diluted HEPES buffer (50 mM, pH 8.5) under 365 nm UV light for 20 minutes, washed twice with HEPES buffer and incubated overnight at 4°C with 100 µg/ml PureCol® Type I Collagen Solution Bovine (Advanced Biomatrix #5005).

### 2D culture

Experiments with HeLa and HeLa_PKD3^EN^-mGFP cells were performed using 2D culture. Cells were cultured on tissue-culture plates or on glass coverslips coated with 2.5 µg/ml Collagen R Solution (Serva #47254) for 1 h at 37°C or overnight at 4°C. When transient transfection was performed, cells were fixed or harvested 72 h post-transfection.

### 3D-on top culture

Unless indicated, all experiments with TNBC cell lines were performed using 3D-on top culture. Transient transfection (when applicable) was performed 16-24 hours prior harvesting and re-seeding cells on hydrogels.

This protocol was adapted from the one described in (Fattet et al., 2020). Collagen-coated PAA hydrogels were prepared as described above, rinsed with PBS and UV-sterilized for 20 minutes. TNBC cell lines were collected by trypsinization, and the appropriate cell count was resuspended in ice-cold DMEM-high glucose supplemented with 10% FBS, 1% Penicillin/Streptomycin and 2% Matrigel® Matrix (Corning #356231), plated on hydrogels and incubated for 5 days.

### Sphere formation assay

TNBC cells were seeded on 5 kPa PAA hydrogels using the protocol for 3D-on top described above. After 5 days, cells were collected by trypsinization, singularized using a 27G needle, resuspended in sphere formation medium (DMEM/F12 supplemented with 10 μg/ml insulin (Sigma-Aldrich #I6634), 20 ng/ml EGF (R&D Systems #236-EG-200), 1 μg/ml hydrocortisone (Sigma-Aldrich #HO888), 1x B27-supplement (Gibco #17504044) and 1% Penicillin/Streptomycin), seeded onto Poly(2-hydroxyethyl methacrylate)-(pHEMA, Sigma-Aldrich)-coated plates in triplicates and incubated for 5 days. 3 x 10^3^ cells were seeded in 1.5 ml/well in a 12-well plate. Spheres were imaged on day 5 and analysed using a semiautomated macro in *Fiji*. First, the spheres were outlined using the function *Sharpen* and *Find edges*, then thresholded and converted to binary images. Holes and borders were closed using 3 iterations of the functions *Erode* and *Dilate*, and the area was quantified using *Analyze particles*. Stemness was evaluated by the sphere formation efficiency (SFE, number of spheres formed per 1000 cells seeded). Only spheres > 2500 µm^2^ were considered.

### Western blot

Whole-cell lysates were prepared in RIPA buffer (50 mM Tris pH 7.4, 150 mM NaCl, 1 mM EDTA, 1% NP40, 0.25% sodium deoxycholate, 0.1% SDS) with phosphatase (PhosSTOP™, Roche) and protease inhibitors (cOmplete™ EDTA-free, Roche), clarified by centrifugation at 13000 rpm for 15 min and quantified using the DC Protein Assay Kit (Bio-Rad #5000112). Equal amounts of protein were loaded, separated by SDS-page (NuPAGE, Invitrogen) and transferred into a nitrocellulose membrane using an Iblot gel transfer device and iBlot Transfer Stacks (both from Invitrogen). The membranes were incubated with Blocking solution (Roche #11096176001) in PBS containing 0.05% Tween-20 and the antibodies as indicated, diluted in Blocking solution. Acquisition was done using an Amersham™ Imager 600 device (Cytiva) and the respective band intensities were quantified in *Fiji* using the raw images and normalised to the loading control (α-tubulin or GAPDH) and to the average intensity for each independent experiment. Figures show a representative membrane after auto-contrast adjustment. All original blots are available as DataSource.

### GST-RILP production and pull-down

The purchased glycerol stock from *E.coli* transformed with the plasmid containing the nucleotides 658-897 of the murine Rab-interacting lysosomal protein (RILP) protein, fused to the C-terminus of GST in the pGEX 4T-3 vector (Addgene #79149, Romero Rosales et al., 2009), was inoculated in 5 ml of LB medium with Ampicillin. Then, 1 ml of the overnight culture was used to inoculate 250 ml of LB medium with Ampicillin and grown at 37 °C to an optical density (OD) of 0.65. Then, IPTG was added to a final concentration of 0.5 mM to induce protein production, and the culture was incubated for additional 4 h at 30 °C, then spun down at 5000 rpm for 10 min, after which the supernatant was discarded, and the bacteria pellet frozen at 20°C. The bacteria were lysed in 5 ml of cold bacteria lysis buffer (25 mM Tris-HCl pH 7.4, 1M NaCl, 0.5 mM EDTA, 0.1% Triton-X, 1 mM DTT) with complete protease inhibitors, and sonicated in 3 cycles of 30 s using a Sonoplus ultrasonic homogeniser. Then the lysates were cleared by centrifugation for 10 min at 13000 rpm. Subsequently, 5 ml of lysis buffer were added to the cleared lysate prior incubation with 300 µl of a pre-equilibrated 50% slurry of glutathione-Sepharose 4B beads for 1 h at 4°C with continuous rotation. The beads were centrifuged at 1500 rpm for 3 min, washed twice with lysis buffer and resuspended as a 50% slurry in lysis buffer. An estimation of the concentration as well as verification of the correct binding was performed by SDS-PAGE and subsequent Coomassie staining.

For the pull-down assays, whole-cell lysates were prepared using 500 µl of lysis buffer (10 mM HEPES, 150 mM NaCl, 5 mM MgCl_2_, 5% Glycerol, 0.1% NP-40, 1mM DTT) with phosphatase (PhosSTOP™, Roche) and protease inhibitors (cOmplete™ EDTA-free, Roche), and passed through a 25G needle ten times to solubilize membrane-bound proteins. After 5 min incubation on ice, lysates were clarified by centrifugation at 13000 rpm for 15 min. 50 µl were kept as input and the rest was incubated in a final volume of 1 ml lysis buffer with 10 µl of a 50% slurry GST-RILP beads coupled to glutathione-Sepharose 4B beads, for 1 h at 4 °C with continuous rotation, spun down and washed thrice with washing buffer (10 mM HEPES, 150 mM NaCl, 5 mM MgCl_2_, 5% Glycerol, 1mM DTT). The supernatant was removed completely, and the beads were eluted by boiling in 5x Laemmli-sample buffer for 10 min at 96 °C.

### Quantitative Real-Time PCR (qRT-PCR) Analysis

RNA was isolated from cells using NucleoSpin RNA kit (Macherey-Nagel, #740955). 80 ng per sample were loaded and one-step RT-PCR reaction was performed in triplicates using Power SYBR® Green RNA-to-C_T_^TM^ 1-Step Kit (Applied Biosystems, #4389986). The following primers were used: *Axin2* (F:agtgtctctacctcatttccc, R:ccctctctctcttcatcctc), *PRKD3* (QuantiTect Primer Assay, Hs_PRKD3_1_SG, NM_005813) and *RPLP0* (F:ctctgcattctcgcttcctggag, R: cagatggatcagccaagaagg). Analysis was performed using the CFX96 Touch Real-Time PCR Detection System (Bio-RAD). mRNA expression values were generated using ΔCt values normalised to *RPLP0*.

### Immunofluorescence and confocal imaging

Cells were washed with PBS, fixed with 4% paraformaldehyde (PFA) for 20 minutes at room temperature, permeabilized with PBS-0.1% Triton X-100 for 5 minutes, and incubated with blocking buffer (PBS-5% FBS). Samples were incubated with primary antibodies overnight in blocking buffer, washed 3 times with PBS, incubated with secondary antibodies with or without 1 µg/ml DAPI for 1 hour at room temperature, washed 3 times with PBS and embedded in ProLong™ Gold Antifade Mountant (Invitrogen #P36930). All samples were analysed at room temperature using a confocal laser scanning microscope (LSM 980 Airyscan 2) equipped with a LD LCI Plan Apochromat 40x/1.2 Imm Corr DIC M27 water-immersion objective for cells cultured on hydrogels, or with a Plan-Apochromat 63x/1.40 DIC M27 for cells cultured on glass coverslips (both from Carl Zeiss). The respective laser excitation wavelengths and emission detection intervals were used as follows: 653 nm and 574–720 nm for Alexa647+; 557 nm and 380–608 nm for Alexa-Fluor546 or Alexa-Fluor555+; 493 nm and 499–548 nm for Alexa-Fluor-488, Alexa-Fluor488+ or GFP and 353 nm and 422– 477 nm for DAPI. Images were acquired in Z-stacks of 250 nm intervals throughout the cell and processed using *ZEN blue* software (Carl Zeiss). Each set of samples was acquired using the same laser settings. For analysis, a single middle Z-section or maximum intensity projections were used as indicated in the figure legends. Images were further analysed in *Fiji* or *ZEN lite* software as described below.

### Endolysosomal acidification

Cells cultured on 5 kPa hydrogels were incubated with 2 µM LysoSensor™ Yellow/Blue DND-160 (Thermo Fisher Scientific #L7545) for 5 minutes prior washing, fixation and mounting as described above. To detect variances in the more acidic lysosomes, the probe was excited at 405 nm and the emission spectra was detected at 451, 459, 468, 477, 486, 495, 504, 512, 521, 530, 539, 556, 565, 574, 583 and 592 nm of a single middle section. For quantification, images were randomized and the mean fluorescence intensity (MFI) of at least 5 vesicles per cell was evaluated using *ZEN lite*. The background was subtracted using the signal in the cell nucleus. An average fluorescence per cell normalized to the average intensity at 451 nm was used to create histograms for each individual experiment, and statistical analysis was performed using the area under the curve (AUC), in three independent experiments.

### Image analysis

The analysis of immunofluorescence images was performed using *Fiji* or *ZEN lite* softwares. The macros used for quantification in *Fiji* are available at https://github.com/elena-gutierrez-galindo/Gutierrez-galindo-et-al.

### 2D vs 3D-on top PKD3-GFP vesicles (Figure 1B)

Maximum intensity projections (MIPs) were used for this quantification. The cell outline was drawn using the *Freehand selection* tool in the GFP channel with increased brightness, and the background was cleared. For each cell, the GFP and pPKD channels were split and a threshold was set for each channel to segment the respective vesicles and converted to a binary image. Then, using the function *Colocalization threshold*, the area between the two channels was segmented and further thresholded into a new binary image, and the pPKD-PKD3 vesicle area was obtained with the *Analyze particles* tool.

### PKD3EN-mGFP vesicles (Figure 1G, S1E)

MIPs were used for this quantification. The cell outline was drawn, and the background was cleared as described. For each cell, the GFP signal was smoothed using the *Gaussian blur* filter and the PKD3^EN^-mGFP vesicles were detected and counted using the *Find maxima* tool. Of note, this function always finds at least one local maxima, so PKD3^EN^-mGFP count per cell is always 1, even when no clear vesicles are visible.

### Rab7 intensity and distribution

MIPs were used for this quantification. The cell outline was drawn, and the background was cleared as described. The Rab7 MFI in the whole cell was obtained. Then, the signal noise was reduced using the function *Subtract background*, and the Rab7 vesicles were identified and saved as ROIs using the plugin *TrackMate* (Ershov et al., 2022). Finally, the ROIs were opened for each cell, the cytosolic background was subtracted, and the same threshold was set for all experiments to segment the Rab7-positive vesicles, then they were converted to binary images and measured using the *Analyze particles* tool.

### Colocalization of Rab7 with Lamp1/CD63 or Vps35

The cell outline was drawn, and the background was cleared as described. The Lamp1, CD63 or Vps35 vesicles were segmented for each individual Z-section and saved as ROIs. Colocalization of Rab7 with these proteins was quantified as the MFI of Rab7 measured for these ROIs and plotted as average intensity per cell.

### Statistical analysis

Data are presented as mean ± SD. Statistical analysis was performed using the software *GraphPad Prism 10*. Significance between two groups was assessed by t-test using two-tailed unpaired analysis. In studies combining knockdown and treatment, p-values were obtained by two-way ANOVA with Šídák’s multiple comparisons test, between control and knockdown for each treatment or between treatments for each siRNA/shRNA group. In studies with different treatments, a one-way ANOVA with Dunn’s multiple comparisons test was used. For confocal microscopy experiments, small datapoints on the superplot represent the average per cell, and the bigger datapoints represent the mean value per experiment, used for statistical analysis. Unless indicated, all experiments were performed at least thrice. Symbols shape in graphs represent independent experiments. The sample size (N), replicates (n) and statistical test are indicated in the figure legends.

## Key resources table

| Reagent | Source | Identifier |  | Additional information |
| --- | --- | --- | --- | --- |
| Antibodies and fluorescent dyes |  |  |  |  |
| α-Tubulin | Merk-Millipore | Cat #05-829 | RRID:AB_310035 | WB 1:10000 |
| Cathepsin D | Cell Signaling Technologies | Cat #74089 |  | WB 1:1000 |
| CD63 | Sigma-Aldrich | Cat #SAB4700215 | RRID:AB_10895913 | IF 1:1000 |
| GAPDH | Sigma-Aldrich | Cat #G9545 | RRID:AB_796208 | WB 1:2000 |
| Lamp1 | Cell Signaling Technologies | Cat #9091 | RRID:AB_2687579 | IF 1:200 |
| Lamp1 | Cell Signaling Technologies | Cat #15665 | RRID:AB_2798750 | IF 1:200 |
| Non-phospho (Active) β-Catenin | Cell Signaling Technologies | Cat #19807 | RRID:AB_2650576 | WB 1:1000 |
| Phospho-PKD/PKCμ (Ser744/748) | Cell Signaling Technologies | Cat #2054 | RRID:AB_2172539 | IF 1:100 |
| PKD2 | Cell Signaling Technologies | Cat #8188 | RRID:AB_10829368 | IF 1:100 |
| PKD3/PKCν | Cell Signaling Technologies | Cat #5655 | RRID:AB_10695917 | WB 1:1000 |
| Rab7 | Cell Signaling Technologies | Cat #9367 | RRID:AB_1904103 | IF 1:50; WB 1:1000 |
| Rab7 | Cell Signaling Technologies | Cat #95746 | RRID:AB_2800252 | IF 1:100 |
| TGN46 | Bio-Rad | Cat #AHP500GT | RRID:AB_2203291 | IF 1:300 |
| Vps35 | Cell Signaling Technologies | Cat #81453 | RRID:AB_2923525 | IF 1:100, WB 1:1000 |
| Donkey Anti-Sheep Alexa Fluor™ 546 | Thermo Fisher Scientific | Cat #A21098 | RRID:AB_2535752 | IF 1:500 |
| Goat anti-Mouse Alexa Fluor™ 488 | Thermo Fisher Scientific | Cat #A11001 | RRID:AB_2534069 | IF 1:500 |
| Goat anti-Mouse Alexa Fluor™ Plus 647 | Thermo Fisher Scientific | Cat #A32728 | RRID:AB_2633277 | IF 1:2000 |
| Goat anti-Rabbit Alexa Fluor™ 546 | Thermo Fisher Scientific | Cat #A11010 | RRID:AB_2534077 | IF 1:500 |
| Goat anti-Rabbit Alexa Fluor™ Plus 488 | Thermo Fisher Scientific | Cat #A32731 | RRID:AB_2633280 | IF 1:2000 |
| Goat anti-Rabbit Alexa Fluor™ Plus 555 | Thermo Fisher Scientific | Cat #A32732 | RRID:AB_2633281 | IF 1:2000 |
| Goat anti-Rabbit Alexa Fluor™ Plus 647 | Thermo Fisher Scientific | Cat #A32733 | RRID:AB_2633282 | IF 1:2000 |
| Peroxidase AffiniPure™<br>Goat Anti-Mouse IgG (H+L) | Jackson ImmunoResearch | Cat #115-035-062 | RRID:AB_2338504 | WB 1:10000 |
| Peroxidase AffiniPure™<br>Goat Anti-Rabbit IgG (H+L) | Jackson ImmunoResearch | Cat #111-035-144 | RRID:AB_2307391 | WB 1:10000 |
| LysoTracker DND-99 Red | Invitrogen | #L7528 |  |  |
| LysoSensor Yellow/Blue DND-160 | Invitrogen | #L7545 |  |  |
| **Chemicals and reagents** |  |  |  |  |
| CRT0066101 | Tocris Bioscience |  |  |  |
| CHIR99021 | Tocris Bioscience |  |  |  |
| **Cell lines** |  |  |  |  |
| MDA-MB-231 |  |  | RRID:CVCL_0062 |  |
| BT549 |  |  | RRID:CVCL_1092 |  |
| HeLa |  |  | RRID:CVCL_0030 |  |
| **Software** |  |  |  |  |
| Fiji |  |  | RRID:SCR_002285 |  |
| GraphPad Prism |  |  | RRID:SCR_002798 | Version 10 |
| Zeiss Zen Lite | Carl Zeiss |  | RRID:SCR_023747 |  |
| Zeiss ZEN (blue edition) | Carl Zeiss |  | RRID:SCR_013672 |  |

---

**Figure S1.**
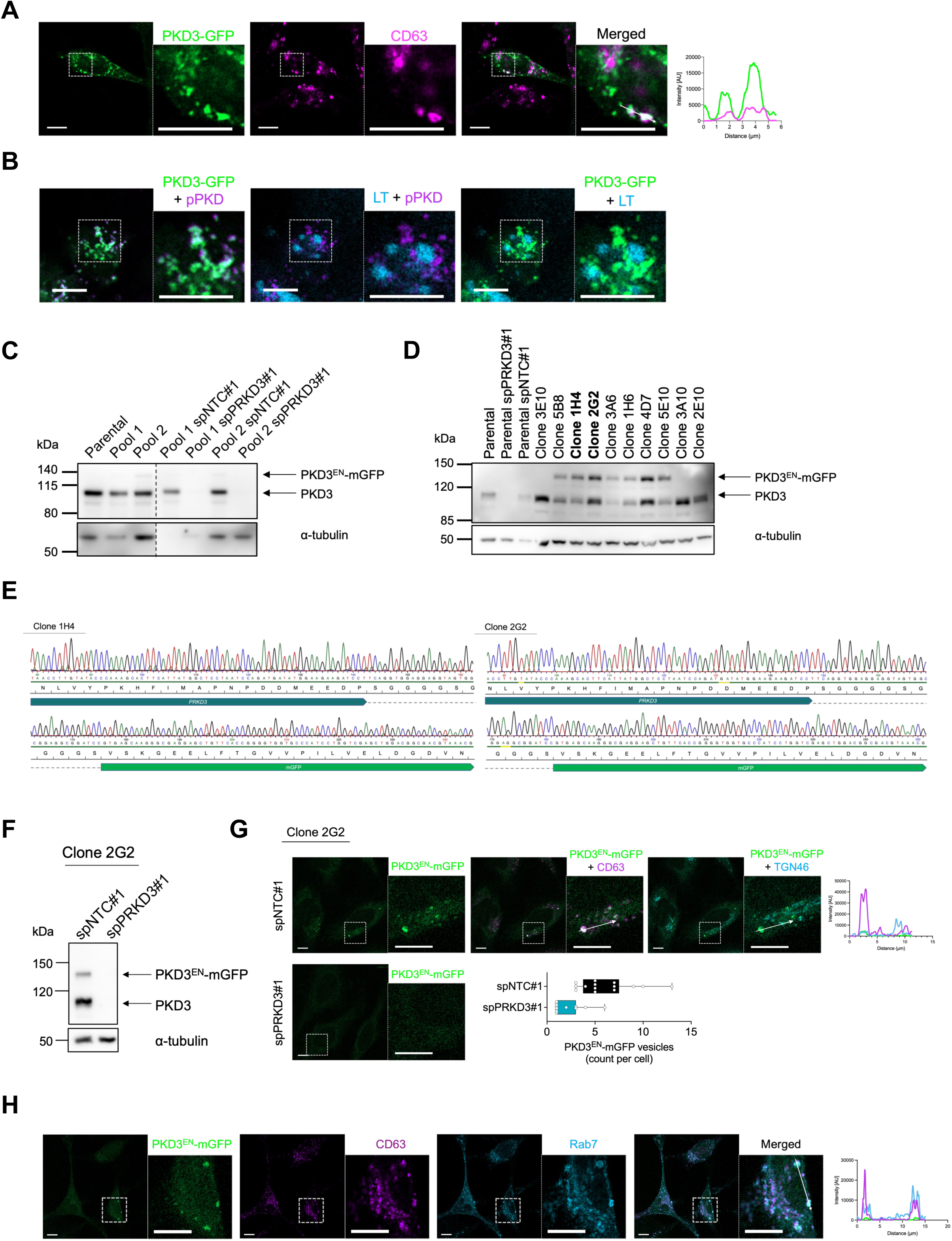
**A-B)** MDA-MB-231_PKD3-GFP cells were seeded on 5 kPa PAA hydrogels, incubated with LysoTracker (in B, blue), fixed and stained for different markers as indicated: CD63 (magenta) in A) and pPKD (magenta) in B); a middle Z-section is shown; scale bar: 10 µm; the histogram represents the intensity profile in the area marked with a white arrow in the merged image. **C)** Parental HeLa cells and two different pools of HeLa_ PKD3^EN^-mGFP cells were seeded and cells from Pool 2 were transiently transfected with spNTC#1 or spPRKD3#1. Whole cell lysates were subjected to Western blot analysis and probed for PKD3; α-tubulin was used as a loading control. **D)** Parental HeLa cells were seeded and transfected with spNTC#1 or spPRKD3#1. 10 different clones resulting from the limited dilution of Pool 2 of HeLa_ PKD3^EN^-mGFP were seeded. Whole cell lysates were subjected to Western blot analysis and probed for PKD3; α-tubulin was used as a loading control. Positive clones selected for further analysis are labelled in bold. **E)** Sequencing of the genomic DNA from HeLa_PKD3^EN^-mGFP (clones 1H4 and 2G2); the amino acids corresponding to the C-terminus of PKD3, the linker and mGFP are indicated. **F)** Whole cell lysates of HeLa_ PKD3^EN^-mGFP (Clone 2G2) transiently transfected with spNTC#1 or spPRKD3#1 were subjected to Western blot analysis and probed for PKD3; α-tubulin was used as a loading control. **G)** HeLa_ PKD3^EN^-mGFP cells (Clone 2G2) transiently transfected with spNTC#1 or spPRKD3#1 were seeded on glass coverslips and fixed; control samples were stained for CD63 (magenta) and TGN46 (blue); scale bar: 10 µm; the histogram represents the intensity profile in the area marked with a white arrow; the graph represents the number of PKD3^EN^-mGFP positive vesicles per cell; N > 10 cells. **H)** HeLa_ PKD3^EN^-mGFP cells (Clone 1H4) were seeded on glass coverslips, fixed and stained for CD63 (magenta) and Rab7 (blue); scale bar: 10 µm; the histogram represents the intensity profile in the area marked with a white arrow in the merged image.

**Figure S2.**
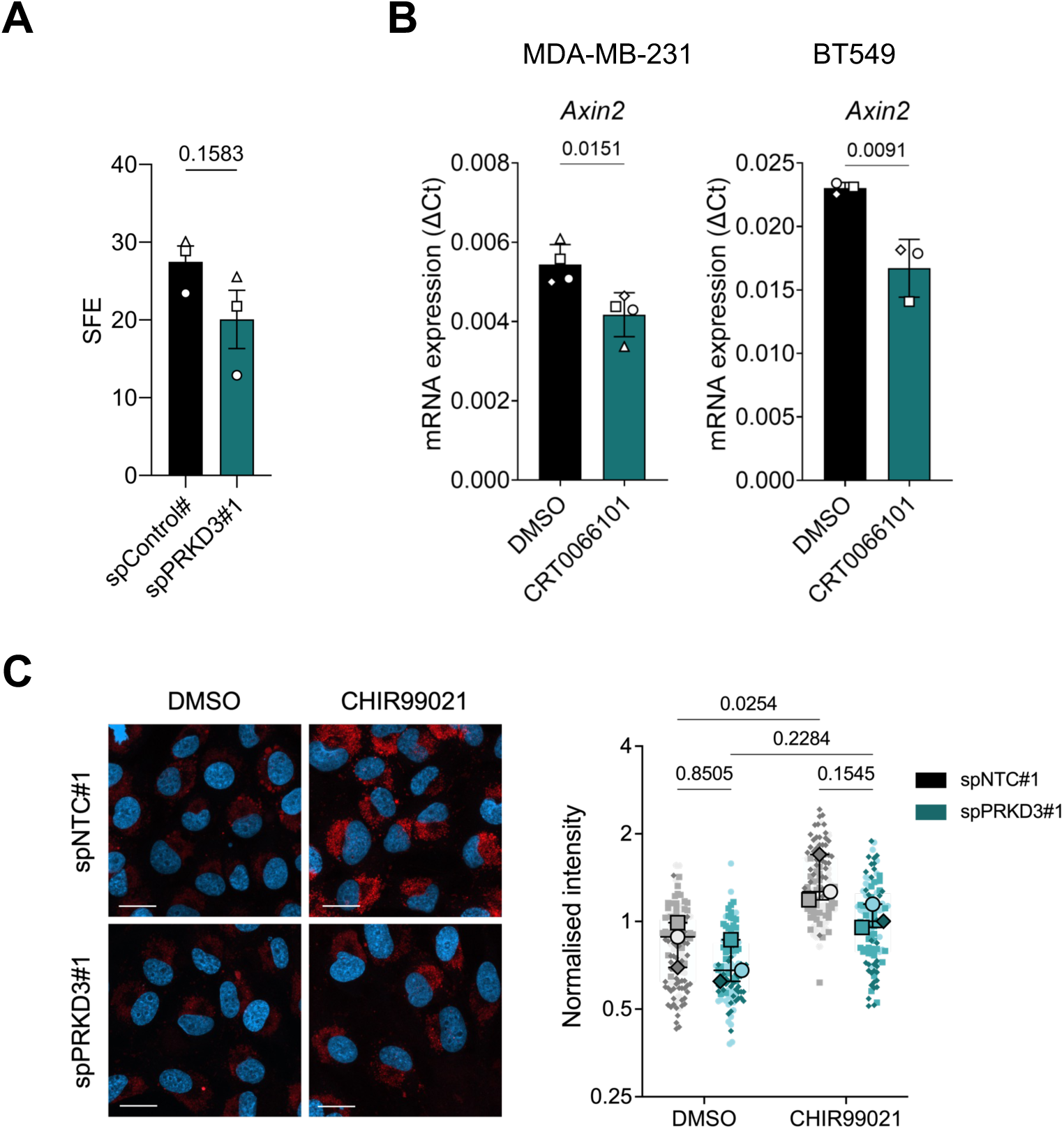
**A)** Quantification of SFA of MDA-MB-231 cells, shown as the sphere formation efficiency (SFE); n=3; statistical comparison by t-test. **B)** qPCR analysis of *Axin1* and *PRKD3* in cells seeded on 5 kPa PAA hydrogels; MDA-MB-231 (left) or BT549 (right) were treated with DMSO or CRT0066101 (1 µM) overnight; n= 3-4; statistical comparison by t-test. **C)** BT549 cells transiently transfected with spNTC#1 or spPRKD3#1 were seeded on 5 kPa PAA hydrogels, treated with DMSO or CHIR99021 (8 nM) for 2 h, incubated with LysoTracker and fixed; scale bar: 20 µm; data in graph represents the MFI, normalized to the average MFI for each independent experiment; statistical comparison by two-way ANOVA with Šídák’s multiple comparisons test; n= 3, N > 35 cells per condition and experiment.

